# Human kinesin-5 KIF11 drives the helical motion of anti-parallel and parallel microtubules around each other

**DOI:** 10.1101/2023.08.01.550848

**Authors:** Laura Meißner, Irene Schüring, Aniruddha Mitra, Stefan Diez

## Abstract

During mitosis, motor proteins and microtubule-associated protein organize the spindle apparatus by cross-linking and sliding microtubules. Kinesin-5 plays a vital role in spindle formation and maintenance, potentially inducing twist in the spindle fibers. The off-axis power stroke of kinesin-5 could generate this twist, but its implications in microtubule organization remain unclear. Here, we investigate 3D microtubule-microtubule sliding mediated by the human kinesin-5, KIF11, and found that the motor caused right-handed rotation of anti-parallel microtubules around each other. The effective sidestepping probability of KIF11 increased with reduced ATP concentration, indicating that forward and sideways stepping of the motor are not strictly coupled. Further, the microtubule-microtubule distance (motor extension) during sliding decreased with increasing sliding velocity. Intriguingly, parallel microtubules cross-linked by KIF11 orbit without forward motion, with nearly full extension. Altering the length of the neck linker increased the forward velocity and pitch of microtubules in anti-parallel overlaps. Taken together, we suggest that helical motion and orbiting of microtubules, driven by KIF11, enable flexible and context-dependent filament organization, as well as torque regulation within the mitotic spindle.

## Introduction

During mitosis, the spindle segregates the chromosomes to the emerging daughter cells. Reliable chromosome segregation is essential to cell survival, as segregation errors may result in chromosome instability and subsequently aneuploidy – a hallmark of several types of cancer. The spindle self-assembles into a metastable structure, with microtubules as basic building blocks. Kinesin and dynein motors organize the microtubules into spindle fibers by cross-linking and sliding them [Walczak et al. 97, Sharp et al. 00]. One of the essential motors in spindle organization is kinesin-5. In prophase, kinesin-5 generates extensile, pushing forces that slide anti-parallel microtubules apart for the segregation of the duplicated centrosomes [Blangy et al. 95, Sharp et al. 99]. During metaphase and anaphase, the sliding activity of kinesin-5 contributes to spindle elongation and its cross-linking activity stabilizes microtubule bundles [Heck et al. 93, Brust-Mascher et al. 09]. Kinesin-5 additionally localizes to the spindle poles, where it bundles and focuses parallel microtubules [Sawin et al. 92, Mann et al. 18]. Inhibition of kinesin-5 impairs pole separation, resulting in monopolar spindles in *Xenopus laevis*, monkey and human cells [Walczak et al. 98, Mayer et al. 99, Kapoor et al. 00]. Kinesin-5 inhibition after spindle assembly causes spindle defects in anaphase B and telophase with effects on positioning of the daughter nuclei in *Drosophila melanogaster* [Sharp et al. 99]. Thus, kinesin-5 is indispensable for the correct functioning of mitosis.

The mechanisms of microtubule organization by kinesin-5 have so far been described by structural and two-dimensional (2D) *in vitro* experimental studies. Kinesin-5 is a bipolar tetramer, with two N-terminal motor domains on each side [Kashina et al. 96]. Each pair of motor domains binds one microtubule, in this way cross-linking the filaments. Upon motor stepping, the microtubules slide apart when their orientation is anti-parallel (microtubule polarity in opposite direction), whereas parallel microtubules do not slide [Kapitein et al. 05]. In addition to forward motion, kinesin-5 displays a sideways stepping component, which results in an orthogonal, sideways motion. When a truncated, dimeric construct of KIF11 was immobilized on a surface, it propelled microtubules forward and simultaneously rotated them in a left-handed manner [Yajima et al. 08]. To explore if sideways components in the power strokes of cross-linking motors can induce helical motion of sliding microtubules around each other, we have recently developed a three-dimensional (3D) motility assay, in which microtubules are suspended on micro-structured polymer ridges [Mitra et al. 18, Bugiel et al. 18]. In this assay the *Drosophila melanogaster* kinesin-14, Ncd, has been shown to drive the right-handed helical motion of short cargo microtubules around the freely suspended sections of immobilized (‘fixed’) microtubules [Mitra et al. 20]. Kinesin-5 is anticipated to show a similar behavior but may display different biophysical and structural properties because of its distinct role in the spindle compared to kinesin-14.

*In vivo*, it has been proposed that the 3D motility of kinesin-5 influences the shape of the spindle, as spindle fibers deform under force. Recently, high resolution imaging of HeLa and RPE1 cells revealed, that the spindle is twisted into a chiral structure. Inhibition of kinesin-5 abolished the twist, whereas overexpression did not change it [Novak et al. 18, Trupinić et al. 22]. This implies that the sideways stepping component of kinesin-5 might be involved in generating rotational forces (torques), which twists the spindle with a specific chirality. In contrast to HeLa cells, spindles of RPE1 cells did not exhibit helicity and only became twisted upon double knockout of kinesin-5 and the dynein targeting factor NuMA, which rendered the spindles fragile [Neahring et al. 21]. Thus, the role of twist is not yet fully understood and, though being a key determinant for spindle shape, the 3D motility behavior of kinesin-5 remains elusive.

Here, we show that human kinesin-5, KIF11, drives the helical motion of anti-parallel microtubules around each other. Interestingly, a similar orbiting motion is also observed for parallel microtubules, though those microtubules are not moving in forward direction. Variation of the critical mechanical element of the motor, the neck linker, increased the helical pitch and velocity of KIF11. This suggests, that KIF11 has evolved as a slowly sliding, fast rotating motor – exhibiting different motility modes dependent on the microtubule orientation.

## Results

To study the 3D motility of microtubules driven by KIF11 we performed 3D sliding assays on micro-structured polymer ridges, with 10 μm wide valleys, separated by 360 nm high and 2 or 5 μm wide ridges (**Fig. 1A, Materials and Methods**). The micro-structures were coated with TAMRA antibodies to suspend long, TAMRA-labeled, ‘fixed’ microtubules. By bridging from ridge to ridge, the lattice of the fixed microtubule is freely accessible over the valley regions. Subsequently, KIF11-EGPF (referred to as KIF11, **Fig. S1, Materials and Methods**) and 1-4 μm long, Atto647N labeled ‘cargo microtubules’ were added in an ADP containing buffer. The sliding process was initiated by adding ATP. Cross-linked cargo and fixed microtubules were tracked using the MATLAB-based software FIESTA [Ruhnow et al. 11] and the perpendicular distance of the center point of the cargo microtubule to the averaged position of the fixed microtubule was measured (referred to as sideways distance). Per our definition, negative sideways distances correspond to movement of the cargo microtubule on the right side of the fixed microtubule (when viewed from the trailing end of the cargo microtubule in the images, **Materials and Methods**).

**Figure 1:**
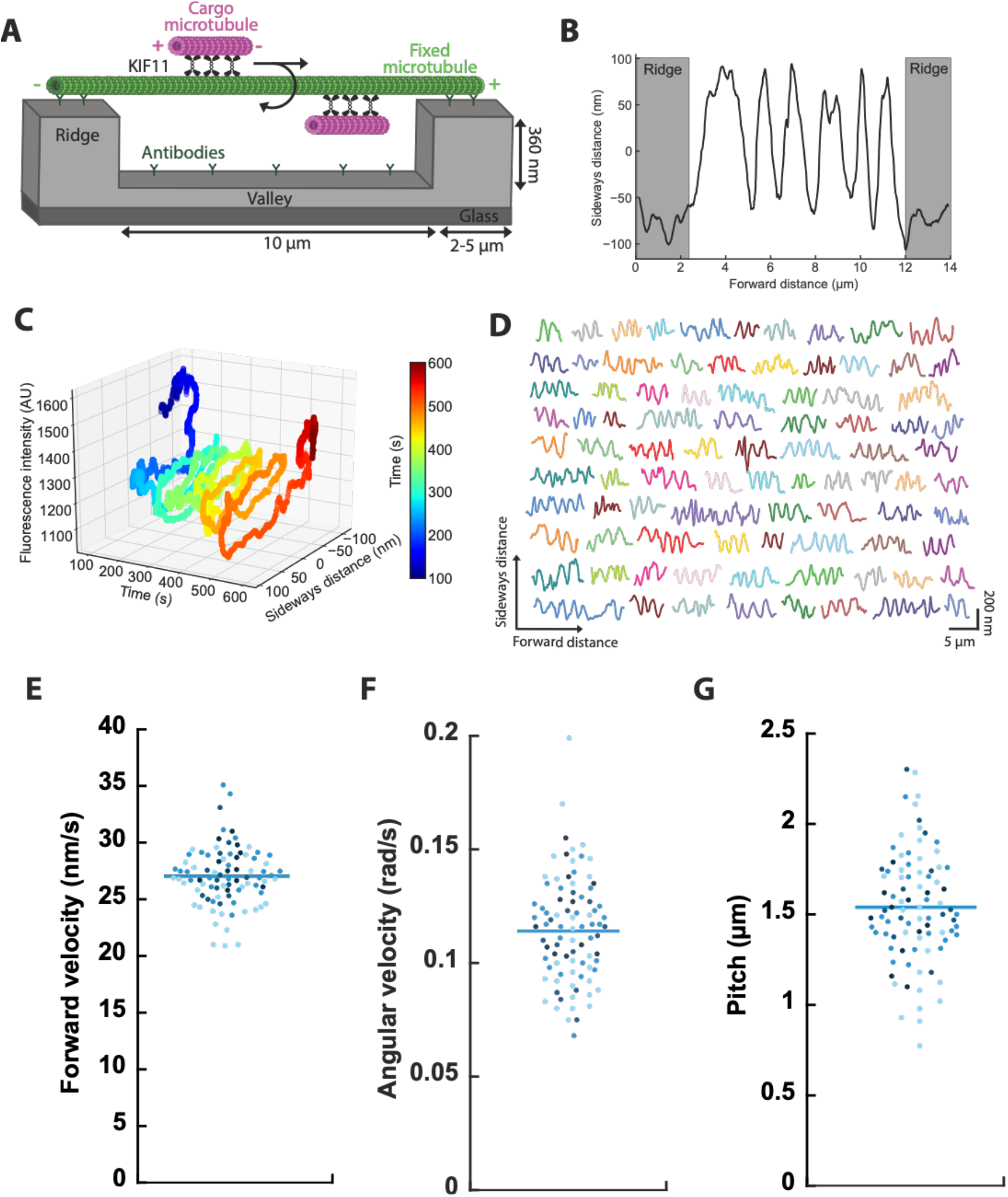
KIF11 drives the right-handed helical motion of anti-parallel cargo microtubules around fixed microtubules. **A**) Setup of the 3D sliding motility assay. Fixed microtubules were suspended on ridge micro-structures, allowing a helical motion of the cargo microtubules driven by KIF11 over the valley regions. **B**) Sideways distance as function of forward distance for an exemplary cargo microtubule. **C**) 3D analysis of cargo microtubule motion. The phase-shifted, periodic changes in the fluorescence intensity and sideways distance of the cargo microtubule are consistent with a right-handed helical motion. **D**) Example tracks of cargo microtubules driven by KIF11 (n = 88). **E - G**) Forward velocity, angular velocity and pitch of the example cargo microtubules from D. Color coding indicates cargo microtubule length: < 1.5 μm (light blue), < 2.2 μm (sky blue), < 2.9 μm (medium blue), >= 2.9 μm (dark blue). Bars indicate mean values.

To analyze the 3D motion, the sideways distance of sliding cargo microtubules was plotted with respect to the travelled forward distance (**Fig. 1B, Supplementary Movie 1**). For the given example, the sideways distance undulated between -100 and 100 nm (showing five full periods) over the valley region, which is reminiscent of a helical motion. Because helical motion was impaired on the ridges, the cargo microtubule was pressed to one side of the fixed microtubule during the first and last 2 μm of forward motion, resulting in a constant sideways distance of around -70 nm. Considering the optical setup of the utilized inverted fluorescence microscope, negative sideways distances on the ridges correspond to a right-handed helical motion (**Materials and Methods**). To confirm the handedness of the helical motion, we lowered the focal plane to the height of the valleys; thus, microtubules were located above the focal plane and the fluorescence intensity increased when the cargo microtubule went from the top of the fixed microtubule to the bottom. A maximum of the fluorescence intensity was then followed by a maximum of the sideways distance with a phase shift of π/2, which confirmed the right-handedness of the helical motion (**Fig. 1C**). In total 88 cargo microtubules displayed similar, robust helical motion over the valley regions (**Fig. 1D**).

The trajectories of the cargo microtubules contain information about their motility parameters. For analysis, only microtubules with at least two full rotations were considered. The motility parameters were calculated for each rotation and averaged for each cargo microtubule (**Materials and Methods**). The forward velocity was 27.0 ± 2.6 nm/s (mean ± standard deviation, n = 88 cargo microtubules, **Fig. 1E**), the angular velocity 0.114 ± 0.023 rad/s (**Fig. 1F**), and the pitch 1.54 ± 0.31 μm (**Fig. 1G**). Neither forward velocity, nor angular velocity, nor pitch correlated with the lengths of the cargo microtubules (**Fig. 1E-G, S2A-C**). We therefore reasoned that the rather large spreads in the motility parameters (in particular angular velocity and pitch) were not due to the length distribution of the cargo microtubules. To investigate if the variability in the motility parameters is inherent or is influenced by minor variations in the fixed microtubule lattice structure, we analyzed eight fixed microtubules, which were traversed by two to six cargo microtubules each. We found that the standard deviations of forward velocity, angular velocity and pitch obtained from cargo microtubules sliding on the same fixed microtubule were similar or higher than the standard deviations of all cargo microtubules on all fixed microtubules: maximum values of 16.6% versus 20.1%, 26.2% versus 9.6%, 30.4% versus 20.2%, respectively (**Fig. S2D** versus **Fig. 1E-G**). This implies that the inherent variability in the motility parameters is not dependent on the variability in the structure of the fixed microtubules.

To test if the sideways motion of cargo microtubules is strictly coupled to their forward motion, we varied the forward velocity by applying different ATP concentrations (1 mM to 25 μM). The forward velocity decreased from 27.0 ± 2.6 nm/s to 5.8 ± 0.7 nm/s and the angular velocity from 0.114 ± 0.023 rad/s to 0.056 ± 0.004 rad/s (**Fig. 2A**). Thereby, the angular velocity showed a weak linear correlation with the forward velocity below 20 nm/s but saturated above 20 nm/s. The pitch decreased with decreasing forward velocity from 1.54 ± 0.31 μm to 0.66 ± 0.09 μm (Pearson correlation coefficient 0.84, **Fig. 2B**). The effective sidestepping probability, calculated as the ratio of sideways movement (in protofilament steps, i.e., in units of 2π/14) to forward displacement (in steps of 8 nm) decreased with increasing velocity from 29.2 to 6.4 (Pearson correlation coefficient -0.86, **Fig. 2C**). This indicates, that the slower a cargo microtubule moves in the longitudinal direction, the more likely it moves sideways in the axial direction. Additionally, we segmented the tracks into ridge and valley parts and grouped them into two types of transitions: from ridge to valley and from valley to ridge. To test for changes in forward velocity at these transitions, we calculated the velocity ratios of both segments (segment 2 divided by segment 1, **Fig. 2D**). We observed that the cargo microtubules did not change their forward velocity at ridge – valley transitions, indicating that suppression of the sideways motion did not affect forward motion. Taken together, these findings show that sideways and forward motion are not strictly coupled.

**Figure 2:**
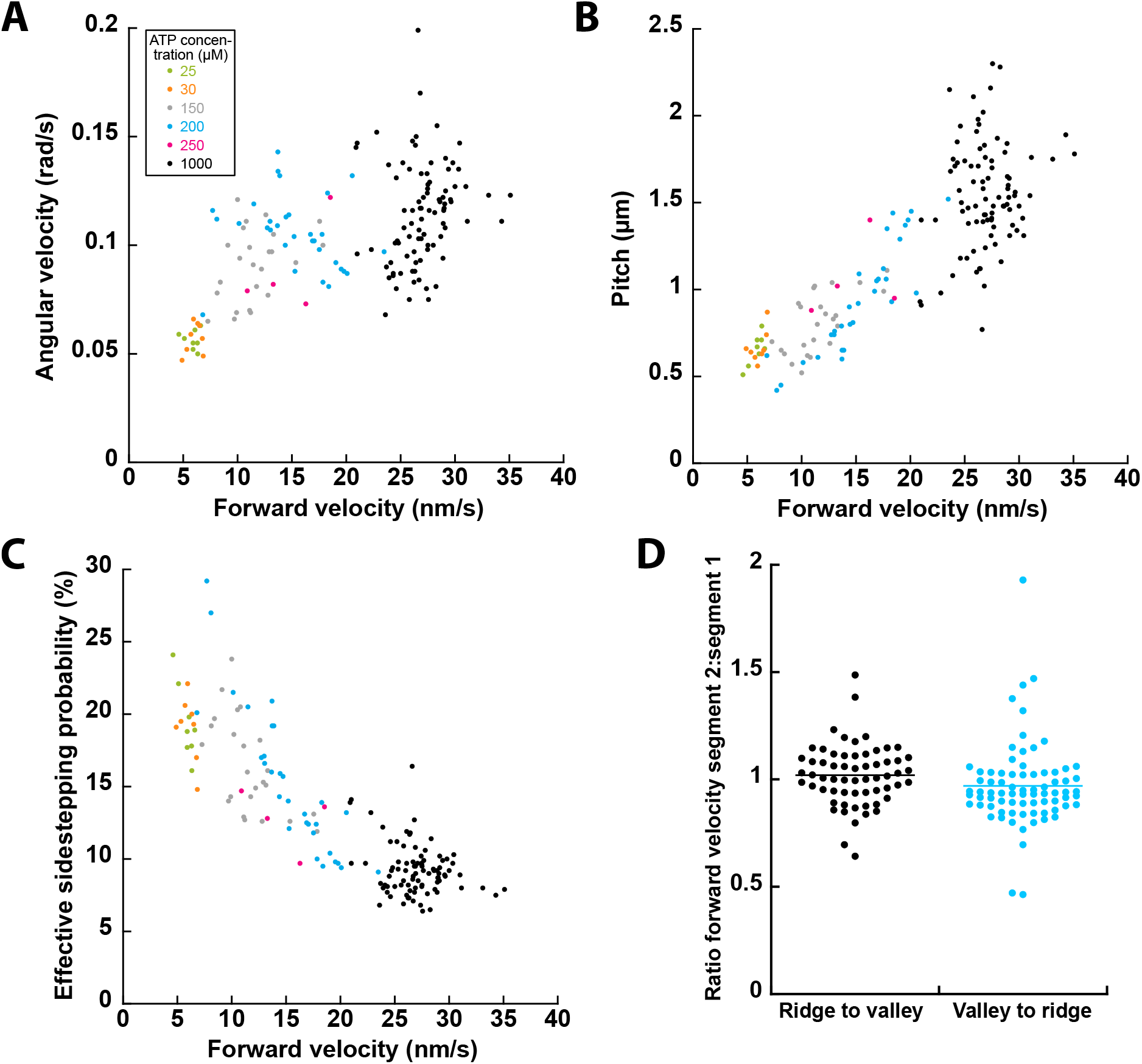
Sideways and forward motion of cargo microtubules driven by KIF11 are not strictly coupled. **A**) Angular velocity, **B**) pitch and **C**) effective sidestepping probability correlated with forward velocity. **D**) The forward velocity of cargo microtubules did neither change significantly at transitions from ridge to valley nor at transitions from valley to ridge.

In our 3D sliding assays, not all cargo microtubules performed a helical motion around the fixed microtubules over valleys. Besides the canonical helical motion (i.e., forward and sideways, **Figs. 1** and **2)**, some cargo microtubules only moved forward (29.4%, forward-only cargo microtubules), while others neither moved forward nor sideways (25.3%, stuck cargo microtubules). For these cargo microtubules we cannot rule out that a helical motion was not detected due to technical reasons (**Materials and Methods**). However, 27.4% of the cargo microtubules did not move forward significantly (forward velocity slower than 5 nm/s) but showed robust orbiting around the fixed microtubules (**Fig. 3A**, sideways-only cargo microtubules, **Fig. S3A-D, Supplementary Movie 2**). Generally, while a cargo microtubule oriented anti-parallel to a fixed microtubule is expected to move forward, parallel microtubules are locked in the longitudinal direction (**Fig. 3B**) [Kapitein et al. 05]. Hence, we conjectured that the helically moving cargo microtubules were anti-parallel to the fixed microtubules, while the sideways-only cargo microtubules were parallel. To test this hypothesis, we employed polarity-labeled cargo microtubules and used the pronounced residence time and accumulation of KIF11-EGFP on the plus ends of the fixed microtubules as markers for the polarity of the fixed microtubules (**Materials and Methods**). We detected five events of helically moving cargo microtubules with four of them in an anti-parallel and one in a parallel orientation (**Fig. S3E, Supplementary Movie 3**). In contrast, from 18 sideways-only cargo microtubules all of them were in a parallel configuration (**Fig. S3F, Supplementary Movie 4**), confirming our hypothesis. Additionally, we observed a number of events where a cargo microtubule did not move forward initially but began to move forward rapidly, after flipping its orientation, with helical motion in anti-parallel orientation and orbiting in parallel orientation.

**Figure 3:**
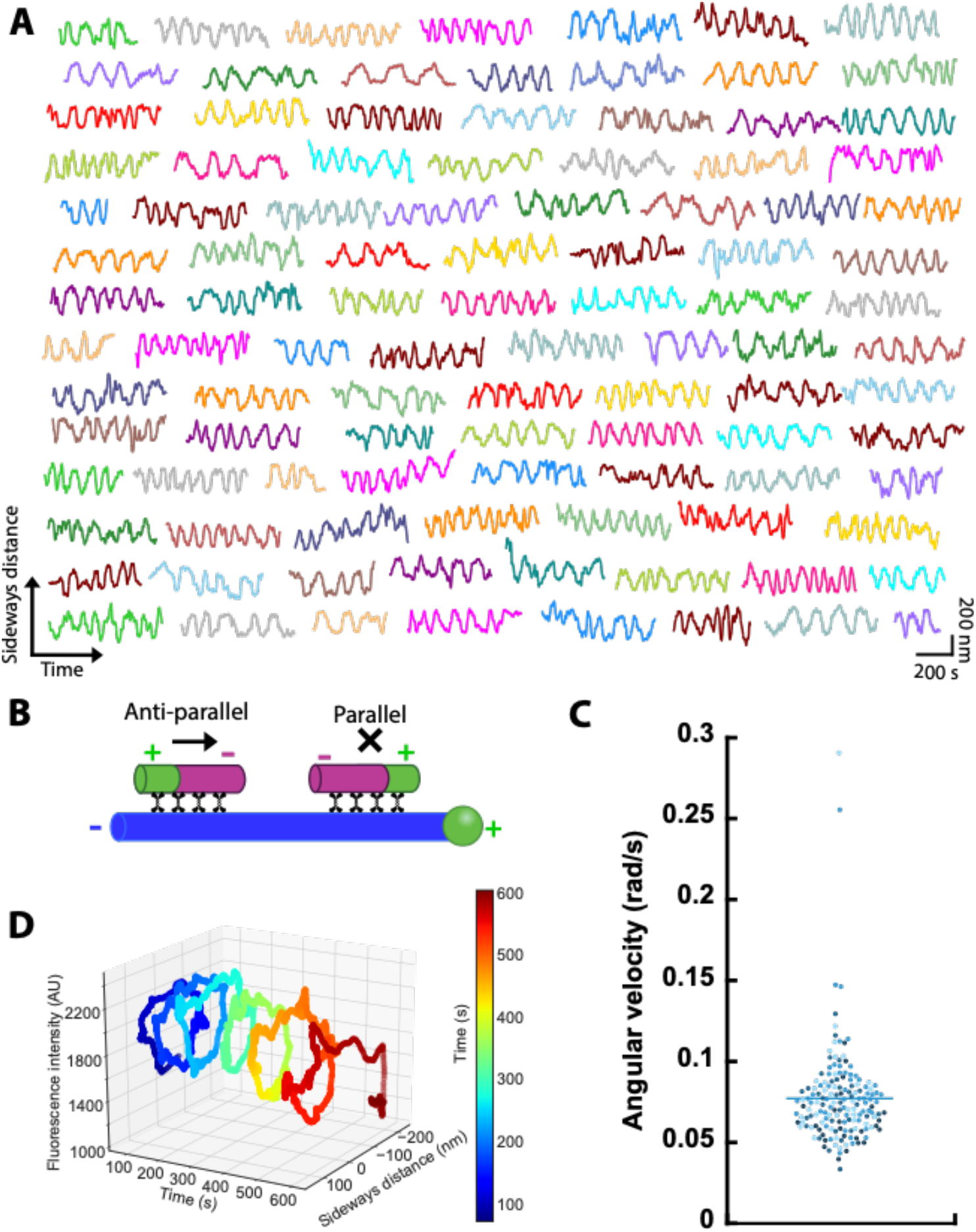
KIF11 orbits parallel cargo microtubules around fixed microtubules without significant forward movement. **A**) Example tracks of cargo microtubules driven by KIF11 (n = 161). **B**) Schematics of microtubule orientation-dependent motility modes of KIF11 in forward direction. **C**) Angular velocities of cargo microtubules moving sideways-only. Color coding indicates cargo microtubule length: < 1.5 μm (light blue), < 2.2 μm (sky blue), < 2.9 μm (medium blue), >= 2.9 μm (dark blue). Bar indicates mean value. **D**) 3D analysis of cargo microtubule motion. The phase-shifted, periodic changes in the fluorescence intensity and sideways distance of the cargo microtubule are consistent with a right-handed orbiting when viewed from the minus end of the cargo microtubule.

Sideways-only cargo microtubules exhibited a 1.5-fold lower angular velocity (0.077 ± 0.029 rad/s) than the forward and sideways moving cargo microtubules (**Fig. 3C**, compared to **Fig. 1F**). Again, no correlation with the lengths of the cargo microtubules was observed. For eight out of nine sideways-only cargo microtubules we confirmed a right-hand orbiting, when viewed from the minus end of the fixed microtubule (**Fig. 3D**). Taken together, KIF11 drives microtubule-microtubule sliding in at least two modes: (i) cargo microtubules which are anti-parallel to a fixed microtubule move forward in a right-handed helical manner around the fixed microtubule with fast angular velocity and (ii) cargo microtubules, which are parallel to the fixed microtubule, orbit in a right-handed manner around the fixed microtubule with slower angular velocity, without forward movement.

An additional parameter that can be estimated from our measurements is the distance between cargo and fixed microtubules during helical motion or orbiting. Under the assumption that the motors bind both microtubules at their shortest distances (i.e., on the protofilaments facing each other), this value represents the extension of the active motors perpendicular to the microtubules (referred hereafter as ‘motor extension’). To this end, we measured the distance between the minima and maxima of the sideways distance and determined the motor extension, as illustrated in **Fig. 4A** and described in **Materials and Methods**). At 1 mM ATP concentration we determined a motor extension of 49.6 ± 9.8 nm (n = 88 overlaps, **Fig. 4B**) which is about 60% of the motor contour length of 79 nm obtained from electron microscopy [Scholey et al. 14]. Upon lowering the velocities by reducing the ATP concentration, the motor extension increased to 64.9 ± 7.1 nm (n = 8). The motor extension in the sideways-only events yielded even higher values of 75.7 ± 14.9 nm (n = 161), close to the motor contour length. Thus, the motors adopt a larger extension the lower the forward velocity and reach almost the maximum extension in absence of forward motion.

**Figure 4:**
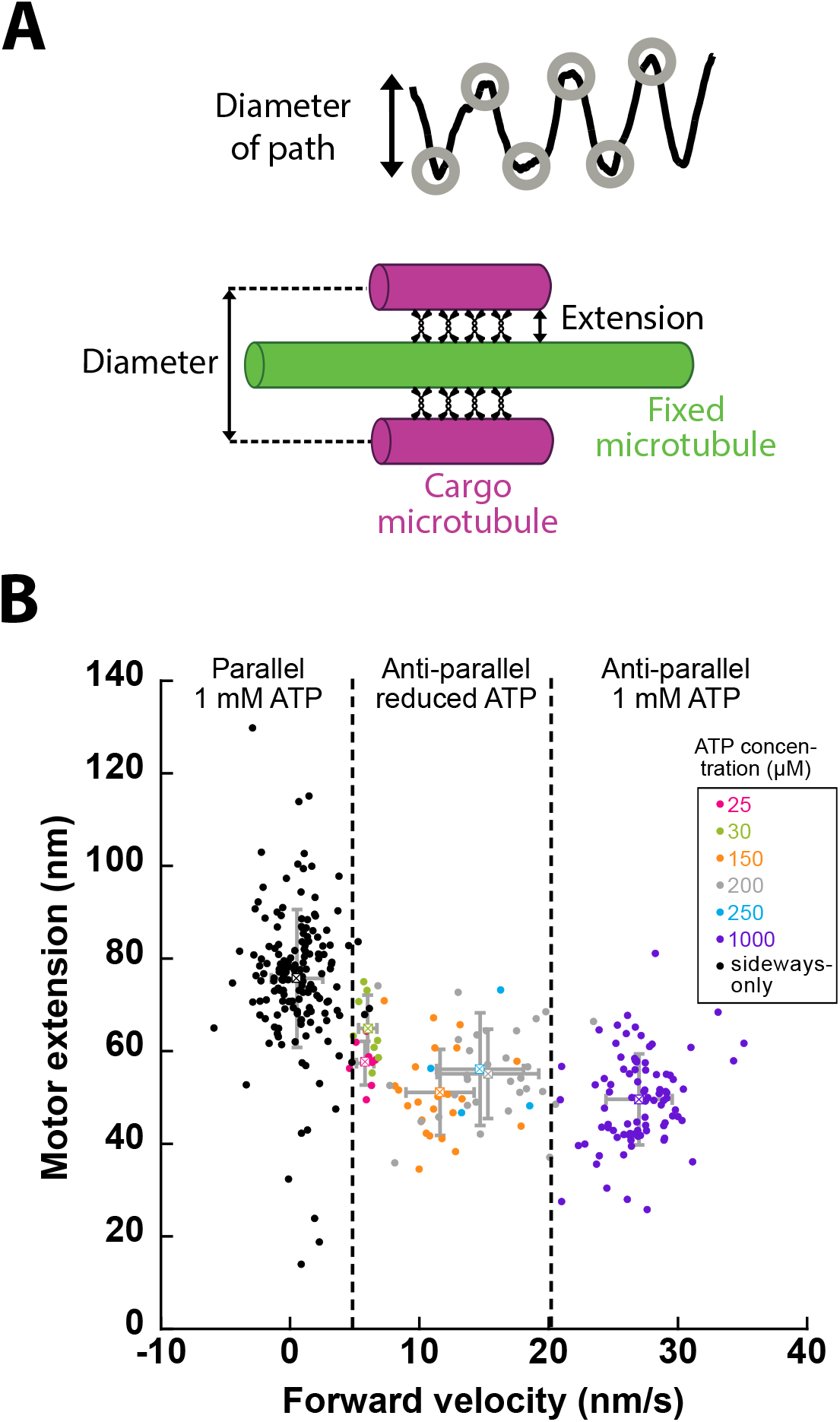
KIF11 motor extension in microtubule overlaps depends on microtubule orientation and forward velocity. **A**) Schematics to estimate the motor extension. **B**) The motor extension of KIF11 correlated with forward velocity (with the lowest extension at highest velocity). Motors in sideways-only events displayed the highest extension.

Finally, we sought to perturb the sidestepping of KIF11 by varying the length of the neck linker domain to identify how it contributes to the sidestepping (NL18-WT, **Fig. 5A**). Previous work found that neck linker length and rigidity affected the sidestepping rate of *Drosophila melanogaster* kinesin-1, KHC. There, an insert of three residues in the same protein changed its rotational pitch (from 8.7 μm of the supertwist to 4.4 μm) [Mitra et al. 18]. Contrary to this, the velocity and force of kinesin-5/kinesin-1 chimeras were independent of the neck linker length [Düselder et al. 12]. We either inserted glycine and serine residues at the C-terminus (GS, NL20 and GSGS, NL22) or removed the C-terminal two or four residues (NL16 and NL14) and tested all constructs in 3D sliding motility assays. The change of the neck linker length did not affect the mobility of cargo microtubules. For the wildtype, 47% of cargo microtubules moved forward (n = 306) and 53% were immobile in forward direction (n = 341). The neck linker mutants displayed a similar fraction of mobile cargo microtubules, between 36 and 58% (all mutants pooled: n = 175). The mean forward velocity drastically increased for all constructs, from 27 ± 3 nm/s of the wildtype to 74 ± 29 nm/s (n = 12), 48 ± 20 nm/s (n = 11), 127 ± 34 nm/s (n = 18), and 136 ± 15 nm/s (n = 7) for NL14, NL16, NL20 and NL22, respectively (**Fig. 5B**). Similarly, the angular velocity of the neck linker mutants doubled to tripled compared to the wildtype from 0.11 to 0.23 – 0.30 rad/s (**Fig. 5C**). The pitch of NL16 (1.2 ± 0.5 μm) was similar to NL18-WT (1.5 ± 0.3 μm) and the pitch of NL14 increased to 2.4 ± 1.5 μm, whereas the pitch of the elongated neck linkers more than doubled to 3.5 ± 2.0 (NL20) and 4.2 ± 1.5 μm (NL22, **Fig. 5D**). Notably, the distribution of both forward velocity and pitch broadened for all neck linker mutants. As previously observed for NL18-WT, the pitch of the neck linker mutants increased with the forward velocity, showing low pitches at low forward velocities (**Fig. 5E**). Similar to NL18-WT, we observed a significant fraction of orbiting cargo microtubules (NL14: one event, NL16: five events, NL20: 15 events, NL22: one event). Thus, the neck linker mutants retain the different motility modes but display drastically altered and more variable motility parameters.

**Figure 5:**
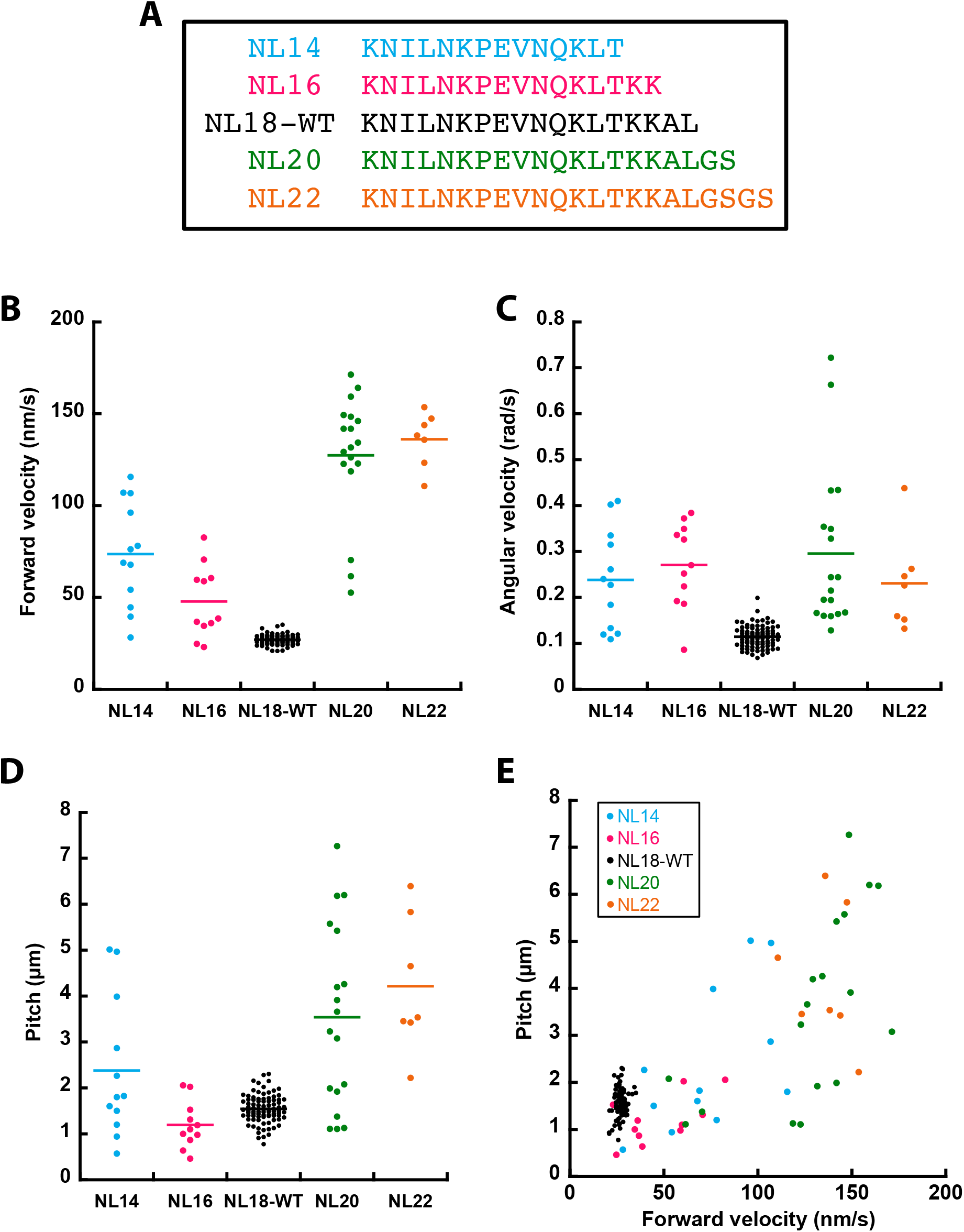
Changing the neck linker length of KIF11 affects the motility parameters. **A**) Overview of neck linker mutant constructs. **B**) Forward velocity, **C**) angular velocity and **D**) pitch of cargo microtubules driven by the neck linker mutants in comparison to wildtype. **E**) The pitch showed a correlation with forward velocity.

## Discussion

Previously, kinesin-5 driven microtubule-microtubule sliding has been studied in 2D motility assays with fixed microtubules fully immobilized on glass surfaces. Such assays have been limited to observing the forward motion of cargo microtubules along fixed microtubules. To investigate the sideways stepping component of motors, 3D setups are required, as demonstrated in recent studies [Brunnbauer et al. 12, Bugiel et al. 18, Can et al. 14, Mitra et al. 18]. Using a 3D setup, where fixed microtubules were elevated on micro-structured polymer ridges, we showed that KIF11 drives the helical motion of cargo microtubules around fixed microtubules. This is the second report of such helical motion for microtubule cross-linking motors after an earlier demonstration for kinesin-14, Ncd [Mitra et al. 20], and the first report for a plus-end directed motor. On the ridges, where the fixed microtubules were immobilized, the sideways motion was sterically blocked and the cargo microtubules were pressed to the right side of the fixed microtubules, reminiscent of a right-handed helical motion.

Our observations are the first example of a right-handed helical motion of a plus-end directed motor protein. Previous reports found that plus-end directed motors (e.g., kinesin-6 and kinesin-8) move along microtubules in a left-handed manner, whereas minus-end directed motors such as kinesin-14 and dynein have been reported to move in a right-handed manner [Maruyama et al. 21, Mitra et al. 18, Nitzsche et al. 16, Can et al. 14, Mitra et al. 20, Walker et al. 90]. We note that the handedness is identical in different assay geometries: left-handed stepping of the motors on microtubules corresponds to a left-handed rotation of microtubules around their long axis in gliding assays and a left-handed helical motion of cargo microtubules around fixed microtubules in 3D sliding assays. The right-handedness in our experiments is contradictory to previous observations of single-headed human Eg5 (KIF11) and truncated (528 N-terminal residues) yeast kinesin-5 Cin8, that rotate microtubules in gliding assays in a left-handed manner [Yajima et al. 08, Yamagishi et al. 21]. It is possible, these conflicting findings arise from differences in protein structure, as KIF11 and Cin8 only share 45% sequence identity (528 N-terminal residues). Additionally, truncation of the motor can also affect its motility, as shown for kinesin-1, which displayed a torque component only in the truncated form [Yajima et al. 05].

The mean pitch of the helical motion of 1.5 ± 0.3 μm in our experiments cannot be related to the supertwist of GMP-CPP grown microtubules, which mainly (96%) consist of 14 protofilaments, resulting in a left-handed supertwist of about 8 μm [Hyman et al. 95, Nitzsche et al. 08, Ray 93]. Thus, the helical motion must arise from a sideways stepping component of KIF11. The pitch is similar to the helical motion driven by Ncd (median pitch of 1.6 μm, [Mitra et al. 20, Yajima et al. 08, Yamagishi et al. 21]), but five times higher than the pitch of single-headed human Eg5 (KIF11) and truncated Cin8 in surface gliding assays (pitches of about 0.3 μm [Mitra et al. 20, Yajima et al. 08, Yamagishi et al. 21]). We believe, the latter difference might be rather attributed to the different motor constructs and different motor properties for different organisms than to the assay geometry – because KIF11 rotated microtubules around their own long axis in surface gliding assays, imaged using fluorescence interference contrast microscopy (FLIC) [Mitra et al. 15], with pitches between 1.1 μm and 4.1 μm (**Fig. S4**). The angular velocity and the pitch of cargo microtubules displayed large spreads of 0.07 – 0.20 rad/s and 0.8 – 2.3 μm, respectively. We could neither attribute these spreads to cargo microtubule length nor forward velocity (Pearson coefficients < 0.3). We also ruled out that the angular velocity and pitch are set by the fixed microtubule, because different cargo microtubules on the same fixed microtubule moved with highly variable motility parameters. Hence, we speculate that the motor density in the microtubule overlaps might vary, depending on the total motor number and length of fixed and cargo microtubules – immobilized and in solution – as well as the history of the fixed microtubule (e.g., motor accumulation). This assumption is further supported by the observation of a higher variability for shorter cargo microtubules, where changes in the density of motors would have a larger relative impact (**Fig. S2A-C)**. Additionally, for the cross-linking motor Ncd it was demonstrated, that the pitch of the rotation of the cargo microtubule around its long axis indeed varied in 2D sliding assays in dependence of motor concentration. Thus, motor density is likely a key factor, which caused variability of motility parameters.

Our observation of similar velocities over ridges and valleys suggests that forward motion is achievable when sideways motion is suppressed, implying that the two motions are not strictly coupled. To validate this hypothesis, we conducted experiments in which we reduced the ATP concentration from 1 mM to 25 μM, resulting in a decrease in forward velocities to 5.8 ± 0.7 nm/s. The effective sidestepping probability then more than doubled, rising from 9.2 ± 1.8% at 1 mM ATP to 19.4 ± 2.6% at 25 μM ATP. These results indicate that at lower ATP concentrations and subsequently lower forward velocities, KIF11 is more inclined to step sideways. This finding further supports the notion that forward and sideways motion are not strictly coupled, as a strict coupling would have resulted in a constant effective sidestepping probability regardless of the forward velocity.

In the case of yeast kinesin-8, Kip3, and *D. melanogaster* kinesin-14, Ncd, two different models have been proposed to explain their sidestepping behavior. Similar to KIF11, Kip3 is a processive motor and exhibits a dependence of rotations on ATP concentration. A model was proposed for Kip3, where the motor spends an extended period in the ATP waiting state [Mitra et al. 18]. During this waiting time, the motor has a stochastic probability to switch from a less-diffusive two-head bound state to a more diffusive one-head bound state. Upon ATP binding, the motor is significantly more likely to sidestep if it is in the one-head bound state. Thus, the longer the motor waits for ATP, the more likely it is to switch to a one-head bound state and sidestep. On the other hand, the power stroke of the non-processive Ncd consists of a primary forward component along with a smaller lateral component [Maruyama et al. 21, Mitra et al. 18, Nitzsche et al. 16, Can et al. 14, Mitra et al. 20, Walker et al. 90]. Since both KIF11 and Kip3 are processive motors, it is possible that a similar stepping model as that of Kip3, incorporating both ATP-dependent and ATP-independent components, may also apply to KIF11.

Besides the helical motion, we observed that KIF11 has at least one other motility mode, in which cargo microtubules orbit around fixed microtubules without forward movement. Polarity-labelling of fixed and cargo microtubules demonstrated, that cargo microtubules which moved sideways and forward were anti-parallel to the fixed microtubules, whereas cargo microtubules which exhibited a sideways-only motion were parallel. This behavior of the forward motion is not surprising because the forward movement of motors is expected to cancel out in parallel microtubules [Kapitein et al. 05, Fink et al. 09]. Helical motion and orbiting, however, arise from the sideways stepping of the motor domains on the fixed microtubule, independent of the orientation of the cargo microtubule (**Fig. 6A**, black and blue arrows). Ncd did not drive the orbiting of cargo microtubules around fixed microtubules in parallel arrangement, although the same geometrical considerations should apply to this motor. We can only speculate that the non-processive movement of individual Ncd motors prevents persistent orbiting. In addition to the helical or orbiting motion of the cargo microtubules around the fixed microtubules, the KIF11 motor domains are expected to rotate the cargo microtubules around their own axes, with different rotation directions for anti-parallel and parallel arrangements (**Fig. 6A**, grey and purple arrows). It will be intriguing to simultaneously monitor this rotation in addition to the helical and orbiting movement in future experiments.

**Figure 6:**
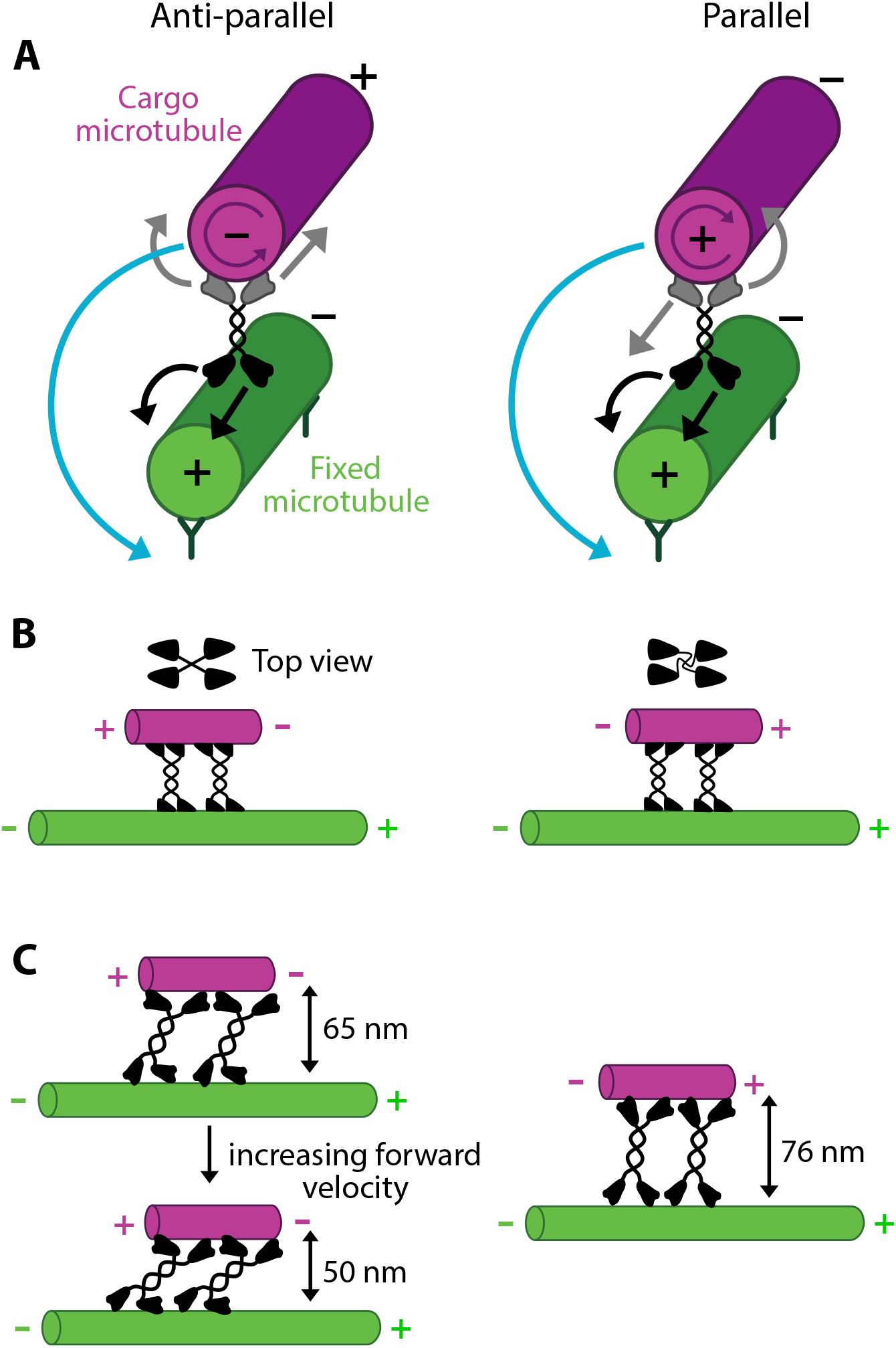
Schematics for helical sliding of anti-parallel microtubules and orbiting for parallel microtubules. **A**) Forward and sideways stepping directions of KIF11 in anti- and parallel microtubule overlaps. Rotational motion occurs in both cases. **B**) In anti-parallel microtubule overlaps, the motor domains of KIF11 interacting with the two microtubules point in opposite directions, whereas they point in the same direction for parallel overlaps (top views of the motor domains indicated). **C**) The extension of KIF11 decreases for increasing sliding velocity of anti-parallel microtubules (left) and adopts an almost fully extended conformation (contour length of KIF11 about 79 nm) in parallel overlaps (right).

Compared to the helical motion of anti-parallel cargo microtubules, the angular velocity of the orbiting cargo microtubules was lower by a factor of 1.5. This decrease could possibly be attributed to the conformation of the motor domain in the overlaps, which is set by the orientation of the tubulin dimers: The motor domains of KIF11 interacting with the two microtubules have to point in opposite directions in the anti-parallel case and in the same direction in the parallel case. On the other hand, structural data of KIF11 showed, that the pairs of motor domains on each side have an offset of 90°. Thus, the motor might have to twist more in the parallel case, which could result in an increased strain in the motor and therefore a less efficient stepping cycle (**Fig. 6B**).

At 1 mM ATP KIF11 adopted an extension of 50 nm perpendicular to the microtubules in anti-parallel overlaps. The extension increased to 65 nm with decreasing forward velocity and was maximal with 76 nm in parallel microtubule overlaps, i.e., in absence of forward motion of the cargo microtubules (**Fig. 6C**). This indicates that the motors inherently favor a large extension, which is decreased during sliding, possibly due to the viscous drag experienced by the cargo microtubule. 50 nm might resemble the minimum extension, in which the motor is able to operate geometrically because stepping might be impeded in more tilted or oblique motor configurations. We hypothesize that motor extension might be another factor for motor functioning and regulation *in vivo* because other microtubule-associated proteins are present in microtubule overlaps. For example, the passive cross-linker PRC1 holds microtubules apart by about 35 nm [Subramanian et al. 10] and kinesin-14 slides microtubules with an extension of 18 – 21 nm [Mitra et al. 20]. As kinesin-5 and kinesin-14 are antagonists, it will be interesting to investigate (i) what is the distance between overlapping microtubules in the presence of both motors, (ii) if kinesin-5 geometrically alters the activity and binding kinetics of kinesin-14, because the latter may not be able to cross-link both microtubules in the presence of kinesin-5, and (iii) if PRC1 influences the extension of the motors during sliding.

We further explored how perturbation of the neck linker affects the 3D motion of KIF11. The neck linker, which connects the motor domain to the stalk, resembles the key mechanical element of the motor, because it transmits force from the motor domain to the stalk and modulates the microtubule-microtubule affinity [Khalil et al. 08, Hwang et al. 08]. In the ADP state, the neck linker points towards the microtubule minus end. Upon ATP binding the neck linker docks and reorients itself towards the microtubule plus end. In this conformation, the neck linker forms the cover neck bundle with the N-terminal extension [Goulet et al. 13]. Previously, it has been shown that kinesins, which track a single protofilament, possess a shorter neck linker (kinesin-1: 14 residues) than sidestepping kinesins (kinesin-2 and kinesin-8: 17 residues, kinesin-5: 18 residues [Shastry et al. 10, Bormuth et al. 12, Hariharan et al. 09]). Changing the neck linker can alter the motor’s stepping in three ways: by disrupting the cover neck bundle, affecting the motor affinity to the microtubule, and influencing the geometry of binding. For the last case, the length of neck linker could determine, how efficiently the lagging head reaches the next tubulin dimer with its step. Both longer and shorter neck linkers increased forward velocity (1.8 to 5-fold), angular velocity (doubling to tripling) and pitch (for three of the four constructs, 1.6 to 2.8-fold). A higher increase in forward than angular velocity resulted in a lower sidestepping rate of the mutants compared to the wildtype, which decreased the pitch of the mutants. Previous work showed that the deletion of the tail increased the forward velocity of KIF11, but came at the expense of less force production [Bodrug et al. 20], which could be the case for the neck linker mutants as well. Thus, we speculate that KIF11 evolved as a slow, potentially high force, motor with a high sidestepping rate.

Applying our insights to the *in vivo* context, i.e., the mitotic spindle, we hypothesize that KIF11 could drive a helical motion of microtubules around each other in the spindle midzone and an orbiting motion at the spindle poles. Right-handed helical motion in the midzone is expected to lead to a right-handed twist of the spindle fibers, which is the opposite of the observed left-handed twist of spindles in HeLa cells [Novak et al. 18]. As Trupinic *et al*. demonstrated, the perturbation of other motors including kinesin-6 MKLP1, kinesin-8 KIF18A and cytoplasmic dynein as well as the microtubule nucleation factor augmin and the passive microtubule cross-linker PRC1 affect the spindle twist in different ways [Trupinić et al. 22]. Thus, other factors besides kinesin-5 might contribute to the twist and other motors could be alternative torque generators. This is in line with the observations of no pronounced twist of EM reconstructed HeLa spindles and less or no twist in non-cancer RPE1 cells [Kiewisz et al. 22, Trupinić et al. 22]. Hence, KIF11 might regulate and balance torques in the spindle, rather than actively generating it.

In summary, our results reveal that KIF11 can drive both helical and orbiting motion of microtubules - i.e., with and without forward motion, respectively - depending on the relative orientation of the microtubules to each other. We found that the distance between the microtubules, and therefore the motor extension, increased with decreasing velocity. Perturbing the neck linker length altered the motility parameters, indicating the KIF11 has evolved for slow and robust motility. We conclude that the helical motion of microtubules driven by KIF11 allows for flexible, context-dependent filament organization and torque regulation in the mitotic spindle.

## Acknowledgments

We thank Corina Bräuer for her technical support and all members of the Diez laboratory for scientific discussions. We acknowledge Régis Lemaitre and the Protein Expression, Purification, and Chromatography Facility of MPI-CBG for generating the baculovirus. We thank the Microstructure Core Facility of the Technology Platform of the Center for Molecular and Cellular Bioengineering at TU Dresden for providing the ridge micro-structures. The Core Facility is supported by the Deutsche Forschungsgemeinschaft (DFG, German Research Foundation) under Germany’s Excellence Strategy – EXC-2068 – 390729961 – Cluster of Excellence Physics of Life of TU Dresden. We would like to acknowledge funding from the German Research Foundation (SFB1027) and the Boehringer Ingelheim Fonds (PhD stipend to L. Meißner).

## Notes

### Competing Interest Statement

The authors have declared no competing interest.

