## Supplementary Info for "Human kinesin-5 KIF11 drives the helical motion of anti-parallel and parallel microtubules around each other"

### Supplementary Figures

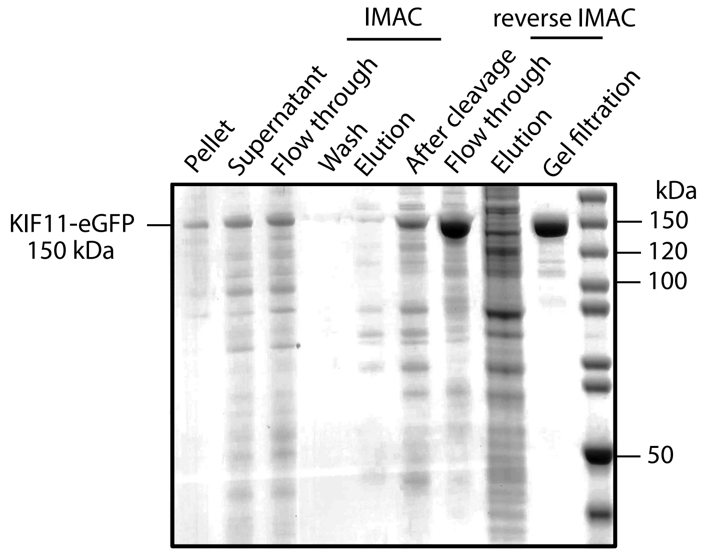


**Figure S1:** SDS gel analysis of KIF11-EGFP purification. KIF11-EGFP was expressed in SF9 cells and samples were taken for the individual purification steps. Lysate was separated in insoluble (pellet) and soluble fraction (supernatant). The supernatant was applied to a HiTrap column (immobilized metal affinity chromatography, IMAC). Proteins without His6 tag did not bind to the column (flow through) and KIF11-EGFP was eluted with imidazole. After tag cleavage, the protein solution was re-applied to the column, did not bind to it (reverse IMAC, flow through) and mostly unspecific proteins were eluted with imidazole. In the last step, the protein solution was subjected to a gel filtration.

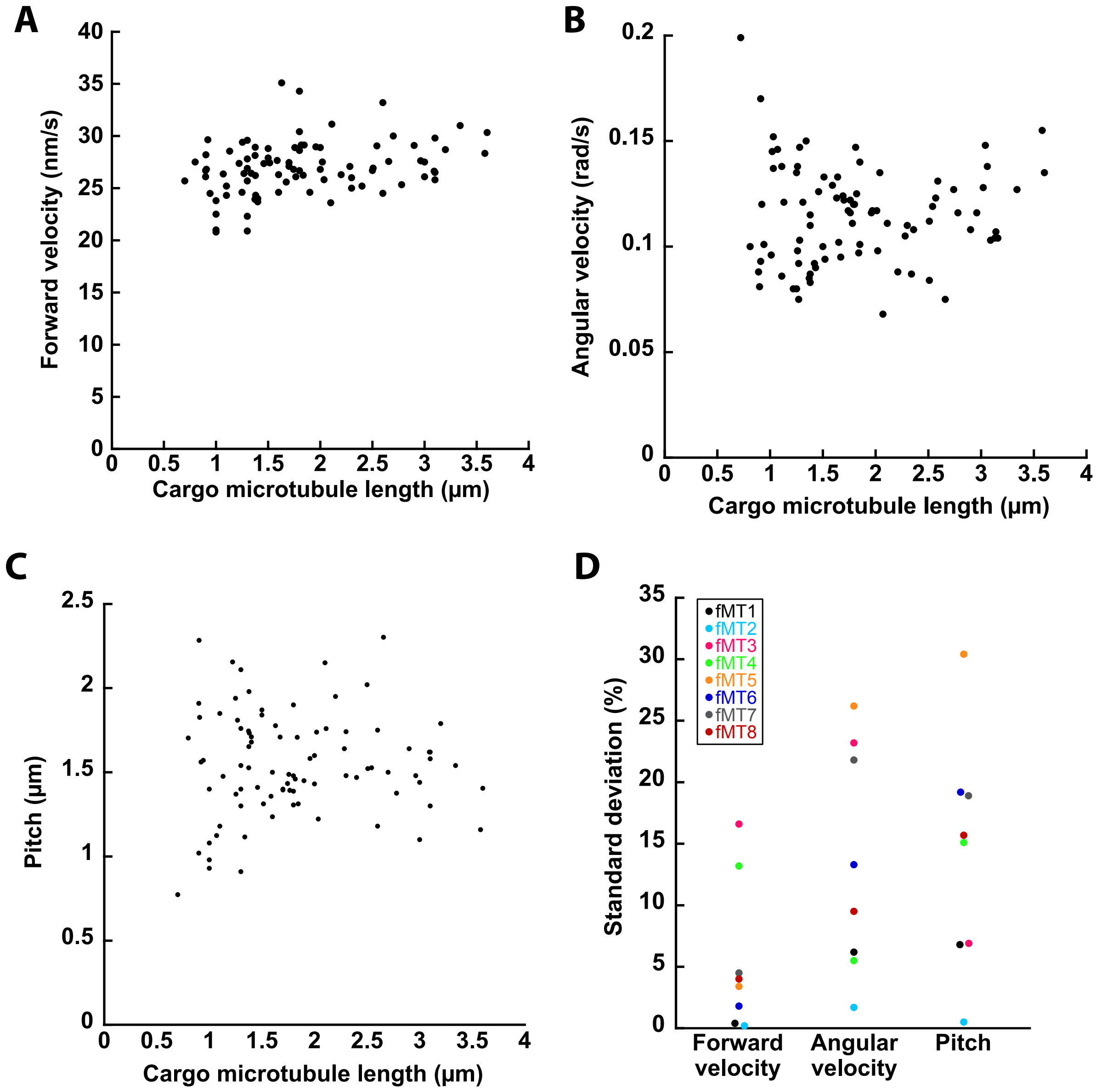


**Figure S2:** Motility parameters as function of cargo microtubule length and particular fixed microtubules. **A**) Forward velocity, **B**) angular velocity, and **C**) pitch as function of cargo microtubule length. **D**) Standard deviations of the motility parameters (in %) for cargo microtubules sliding along particular fixed microtubules (fMT1 - fMT8, color-coded).

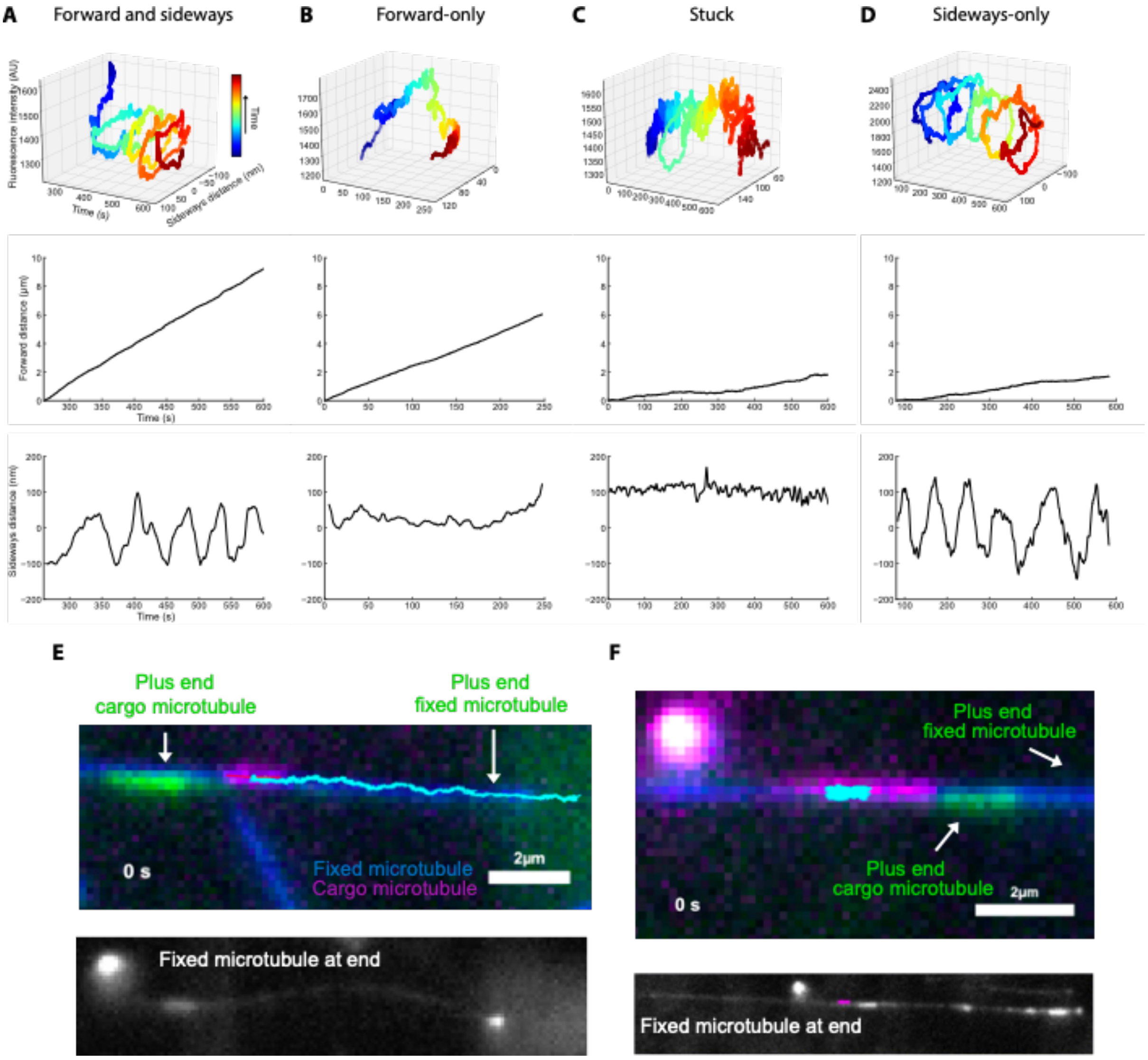


**Figure S3:** Motility modes of KIF11-driven microtubule-microtubule sliding. Cargo microtubules **A**) moved forward and sideways, **B**) moved only forward, **C**) were stuck or **D**) moved only sideways (top row: 3D analysis; middle row: forward displacement; bottom row: sideways displacement). **E**) Forward and sideways event: Color-combined fluorescence micrograph (top) and EGFP signal for determination of microtubule polarity (bottom). Fixed (plus end on right side) and cargo microtubule (plus end on left side) are anti-parallel. **F**) Sideways-only event: Color-combined fluorescence micrograph (top) and EGFP signal for determination of microtubule polarity (bottom). Fixed (plus end on right side) and cargo microtubule (plus end on right side) are parallel.

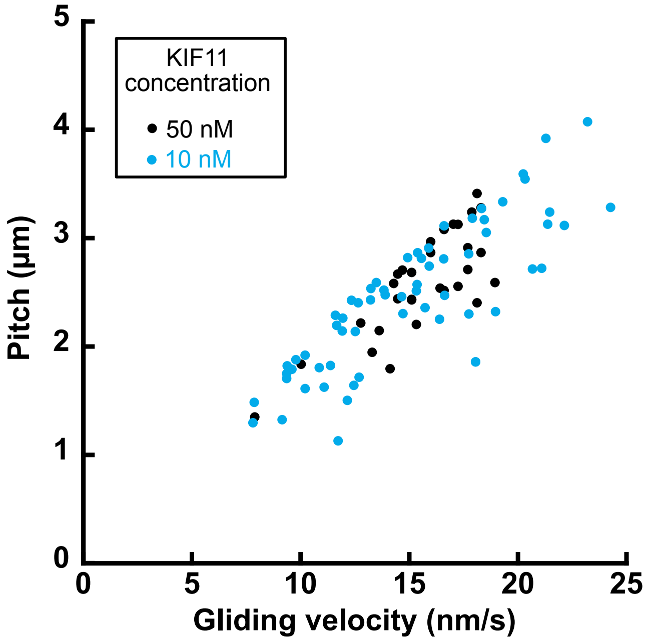


**Figure S4:** Pitches of KIF11-driven microtubule rotations around their own axis in fluorescence interference contrast (FLIC) gliding assays. Pitch correlates with gliding velocity.

### Supplementary Movies

**Supplementary Movie 1:** Example event of a helical motion of a cargo microtubule driven by KIF11. An Atto647N-labeled cargo microtubule (magenta) was sliding along a TAMRA-labeled fixed microtubule (green) suspended over two valleys (10 μm wide) with a ridge (2 μm wide) in between. The tracked filament position of the cargo microtubule is indicated by the red line and the trajectory is indicated by the blue line. Details of this event are also shown in Fig. **1B** and **1C**.

**Supplementary Movie 2:** Example event of an orbiting motion of a cargo microtubule driven by KIF11. An Atto647N-labeled cargo microtubule (magenta) was orbiting around a TAMRA-labeled fixed microtubule (green) suspended over a valley. The tracked filament position of the cargo microtubule is indicated by the red line and the trajectory is indicated by the blue line. Details of this event are also shown in Fig. **3D**.

**Supplementary Movie 3:** Example event of a helical motion of a polarity-labeled anti-parallel cargo microtubule driven by KIF11. An Atto647N-labeled cargo microtubule (magenta) was sliding along a TAMRA-labeled fixed microtubule (blue). The plus end of the cargo microtubule is visible as Atto488-labeled extension (green), imaged before the sliding of the cargo microtubule. The tracked filament position of the cargo microtubule is indicated by the red line and the trajectory is indicated by the cyan line. Details of this event are also shown in Fig. **S3E**.

**Supplementary Movie 4:** Example event of an orbiting motion of a polarity-labeled parallel cargo microtubule driven by KIF11. An Atto647N-labeled cargo microtubule (magenta) was orbiting around a TAMRA-labeled fixed microtubule (blue). The plus end of the cargo microtubule is visible as Atto488-labeled extension (green), imaged before the orbiting of the cargo microtubule. The tracked filament position of the cargo microtubule is indicated by the red line and the trajectory is indicated by the cyan line. Details of this event are also shown in Fig. **S3F**.

**Materials and Methods**

Cloning, gene expression and protein purification. The gene of KIF11 was cut with NotI and AscI and inserted into an OCC vector with C-terminal EGFP and $His_6$-tag, separated by a 3C protease cleavage site. For mutagenesis, the two or four C-terminal residues of the neck linker were removed (NL16 and NL14) or a GS (NL20) or GSGS (NL22) sequence was added at the C-terminus. The above vector was PCR amplified with respective primers: for the shortened constructs the forward primer started after the 3' end of the neck linker and the reverse primer started at the 5' end at amino acid residue 14 or 16 of the neck linker. For the elongated constructs, the forward primer started after the 3' end of the neck linker and bore the respective insert. The reverse primer started at the 5' end of the neck linker.

Viruses were generated using the FlexiBac system[Lemaitre et al. 19]. SF9 cells at 1 million cells per mL were infected with virus (1:100, v/v) and genes were expressed for 96 h at 27 °C and 120 rpm. Cells were centrifuged with 300 g for 10 min at 4 °C. Pellets were resuspended in PBS (1/100$^{th}$ of expression volume) with protease inhibitor, flash frozen in liquid nitrogen and stored at -80 °C. For purification, cell pellets were thawed on ice and resuspended in purification buffer (50 mM $NaH_2PO_4$, 300 mM KCl, 2 mM MgCl2, 1 mM DTT, 0.1 mM ATP, pH 7.5) with protease inhibitor. The lysate was cleared with an ultracentrifuge spin with 40000 rpm for 1 h at 4 °C. The supernatant was filtered through a 0.45 µm filter and loaded on a 1 mL HiTrap column with a superloop. The column was washed with immobilized metal affinity chromatography (IMAC) wash buffer (purification buffer with 20 mM imidazole) and the protein was eluted with IMAC elution buffer (purification buffer with 300 mM imidazole) with an elution gradient. Protein containing fractions were pooled and concentrated with Amicon filters (cutoff 100 kDa). 3C protease was added (1:150, v/v) and the His6 tag was cleaved over night at 4 °C. The protein solution was diluted 6-fold to reduce the imidazole concentration and passed over the HiTrap column again. The protease remained bound to the column with its $His_6$ tag. The flow through was concentrated to 0.5 mL, cleared at 17000 g for 10 min and gel filtered over a Superose6 column with purification buffer. 5% glycerol was added and the protein was flash frozen in liquid nitrogen and stored at -80 °C (**Fig. S1**).

Microtubule polymerization. Tubulin was purified from pig brains according to a standard protocols [Castoldi et al. 03]. Labeling was carried out with labeling kits. Labeled tubulin was mixed in a ratio of 1:3 with unlabeled tubulin for experiments. Fixed and cargo microtubules were grown with tubulin with guanylyl-(α,β)-methylene-diphosphonate (GMP-CPP) and stabilized with taxol. For *fixed microtubules*, 40 µL elongation mix containing 1.25 mM GMP-CPP, 1.25 mM $MgCl_2$ and 4.5 µM TAMRA labeled tubulin in BRB80 (80 mM Pipes at pH 6.9, 1 mM $MgCl_2$, 1 mM EGTA) was incubated on ice for 5 min and then for 30 min at 37 °C. Microtubules were pelleted (17000 g, 15 min) and resuspended in elongation mix (1.25 mM GMP-CPP, 1.25 mM $MgCl_2$ and 0.5 µM TAMRA labeled tubulin in BRB80) and grown for 2 – 3 days at 32 °C. Microtubules were pelleted (17000 g, 8 min) and gently resuspended in BRB80 with 10 µM taxol (BRB80X). They were kept at room temperature for several days for annealing. To grow *cargo microtubules*, a polymerization mix with 1.25 mM GMP-CPP, 1.25 mM $MgCl_2$ and 5 µM Atto647N labeled tubulin was in incubated on ice for 5 min and then for 8 min at 37 °C. Microtubules were pelleted (17000 g, 15 min) and resuspended in BRB80X.

Speckled microtubules (for FLIC experiments, see below) were polymerized based on previous protocols. First, dimly labeled short microtubule seed are grown at a 3 – 10-fold higher tubulin concentration than used for GMP-CPP microtubules. These seeds are added to a solution with bright tubulin at a low concentration to allow for annealing of speckled microtubules. In detail, dimly labeled seeds were polymerized in a mix of 1 mM GMP-CPP, 1.2 mM MgCl2 and 14 µM tubulin (1:90 TAMRA tubulin mixed with unlabeled tubulin) in BRB80. The mix was incubated for 5 min on ice and 25 min at 37°C. Microtubules were pelleted and resuspended in 25 µL BRB80. The elongation mix was assembled (1 mM GMP-CPP, 1 mM MgCl2, 0.8 µM tubulin (1:15 TAMRA tubulin mixed with unlabeled tubulin), 3 µL dimly labeled seeds) and incubated overnight at 37°C. Microtubules were pelleted and resuspended in 50 µL BRB80X.

Fabrication and treatment of ridge structures. Ridge structures were produces and coated with dichlorodimethylsilane as described in Mitra et al. [Mitra et al. 18, Mitra et al. 20].

KIF11 driven microtubule sliding assays. To assemble 6 flow chambers, 7 strips of Nescofilm were placed on the coverslip with structures and covered with a regular coverslip with DDS coating. The Nescofilm was melted on a hot plate. Channels were flushed with: i) 1:60 TetraSpeck bead solution in PBS (v/v, 200 nm) for 2 min, ii) PBS wash, iii) 0.2 mg/mL TAMRA 5G5 antibody solution in PBS for 5 min, iv) 1% F127 (w/v in PBS) solution for at least 1 h, v) 3 washes with BRB80, vi) fixed microtubules in motility buffer (MB-ADP, BRB80 with 10 µM taxol, 200 µg/mL casein, 10 mM DTT, 0.1% (v/v) Tween-20, 20 mM D-glucose, 1 mM ADP), vii) MB-ADP wash, viii) 10 nM KIF11-EGFP for 5 min, ix) MB-ADP wash, x) cargo microtubules, xi) wash with MB-ADP++ (MB-ADP with 200 µg/mL glucose oxidase and 20 µg mL−1 catalase), xii) MB-ATP++ (MB-ADP++ with 1 mM ATP instead of ADP). Where indicated, the final ATP concentration was reduced 250, 200, 150, 30 and 25 µM.

Polarity-labelling of microtubules. For polarity labelling, the plus end of Atto647N labelled cargo microtubules was elongated with Atto488 labelled tubulin. Based on Phelps et al., a maleimidation mix (10 µM Atto488 tubulin, 0.4 mM N-Ethylmaleimide and 1 mM GTP in BRB80) was incubated for 10 min on ice and the reaction was quenched with 20 mM DTT for at least 10 min on ice [Phelps et al. 00]. An elongation mix (4 mM $MgCl_2$, 1 mM GTP, 1 µM Atto488 NEM tubulin and 4 µM Atto488 tubulin in BRB80) was incubated for 5 min on ice. The elongation mix was preheated for 30 s at 37 °C and cargo microtubules were added (tubulin concentration 1 µM). Microtubules were elongated for 5 min at 37 °C, incubated with 5-fold excess BRB80 with 20 µM taxol for 1 min at room temperature, centrifuged at 17000 g for 15 min and resuspended in BRB80X. Polarity-labeled microtubules with Atto488 extensions were also used as fixed microtubules to confirm the pronounced end residency and accumulation of KIF11-EGFP at plus ends.

FLIC assays. The FLIC assay was performed on a 22 mm x 22 mm silanized coverslip and 10 mm x 10 mm silanized SiOx wafers[Mitra et al. 15]. Channels were assembled as for the sliding assays and flushed with i) 0.1 mg/mL anti-GFP antibody solution in PBS for 5 min, ii) 1% F127 (w/v in PBS) solution for at least 1 h, iii) 3 washes with BRB80, iv) 10 – 50 nM KIF11-

EGFP in MB-ATP (ADP replaced with 1 mM ATP in MB-ADP) for 5 min, v) MB-ATP for 5 min, and vi) Speckled microtubule solution for 2 min vii) MB-ATP++.

Optical image acquisition. Optical imaging was performed using an inverted fluorescence microscope (Axio Observer Z1; Carl Zeiss Microscopy GmbH) with a 63× oil immersion 1.46NA objective (Zeiss) in combination with an EMCDD camera (iXon Ultra; Andor Technology) controlled by Metamorph (Molecular Devices Corporation). A LED white light lamp (Sola Light Engine; Lumencor) in combination with a TRITC filterset (ex 520/35, em 585/40, dc 532: all Chroma Technology Corp.), an Atto647N filterset (ex 628/40, em 692/40, dc 635) and a GFP filter set (ex 475/35, em 525/45), corresponding to TAMRA labeled microtubules, Atto647N labeled microtubules and EGFP/mNeonGreen labeled motors, respectively, were used for epifluorescence imaging. The imaging temperature was maintained at 24°C by fitting a custom-made hollow brass ring around the body of the objective and connecting it to a water bath with a cooling/heating unit (F-25-MC Refrigerated/Heating Circulator; JULABO GmbH). The sliding of cargo microtubules was imaged with a 63x oil objective, in the Atto647N channel for 10 min with 10 frames/s with an exposure time of 100 ms. Fixed microtubules were imaged in the TRITC channel for 200 frames at 10 frames/s with exposure time of 100 ms after imaging the cargo microtubules.

Image processing and data analysis. Data was processed and analyzed as described in Mitra et al. [Mitra et al. 20]. Positions of fixed and cargo microtubules as well as TetraSpeck beads (in TRITC and Cy5 channel) were obtained from the MATLAB-based tracking software FIESTA [Ruhnow et al. 11]. The TetraSpeck beads were used to correct the microtubule tracks for drift and color offset. The fixed microtubules were imaged over 200 frames and the tracked positions were averaged to obtain the filament position. The distance of the cargo microtubule’s center point to the fixed microtubule averaged center line was calculated. Negative sideways distances were assigned to cargo microtubules moving in the obtained images on the right side of the fixed microtubule (as viewed from the trailing end of the cargo microtubule which is identical to viewing from the minus end of the fixed microtubule). When no averaged positions of the fixed microtubules were obtainable (e.g., because of interfering signals from small microtubule aggregates close to the fixed microtubule, two fixed microtubules too close together to be resolved or movement of the fixed microtubule between imaging of the cargo and fixed microtubule), the path of the cargo microtubule was averaged over 10 µm and the sideways distance was calculated with respect to the averaged path. For display, cargo microtubule tracks were smoothed over 50 frames.

To obtain the motility parameters with manual computer-aided measurements, the sideways distance was plotted over time and minima and maxima of the rotations were marked. From these points the motility parameters (*forward velocity* = forward distance of the cargo microtubule along the direction of the fixed microtubule per time, *angular velocity* = 2 π divided by the time per rotation, and *pitch* = forward distance travelled per full rotation) were calculated for each rotation and averaged for each cargo microtubule. For ridge-valley comparisons, the forward velocity was calculated the following way: the first and last 50 frames of the time and forward distance of each cargo microtubule track were averaged separately. The fluorescence intensities of the cargo microtubules were obtained from the tracking data of the software FIESTA [Ruhnow et al. 11].

We note, that forward-only microtubules and stuck microtubules could actually be forward-and-sideways microtubules and sideways-only microtubules, respectively, for which the sideways motion was not detected. A technical reason for not detecting the sideways motion could have been an unprecise cargo microtubule tracking due to too jerky or erratic motion. We observed microtubules of all four categories on the same fixed microtubule and thus, can rule out that the fixed microtubule determines the category of the cargo microtubule.

Statistical analysis. Motility parameters were calculated as mean pm standard deviation. Distributions were compared with an ANOVA (analysis of variance) test, $\alpha = 0.05$, followed by a Tukey test to compare two datasets under the assumption of a normal distribution.
